## Supplemental Files for "Unveiling Cell Wall Structure and Echinocandin Response in *Candida auris* and *Candida albicans* via Solid-State NMR"

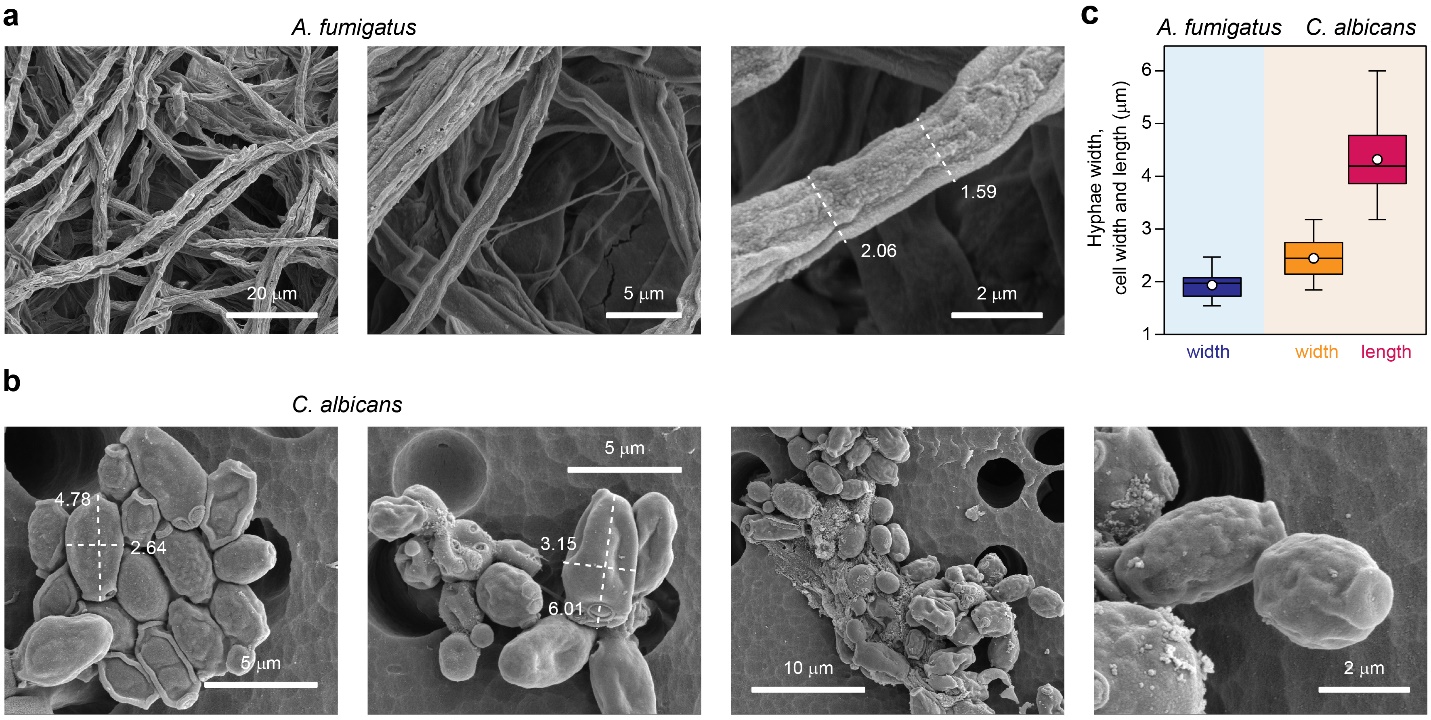

**Supplementary Figure 1. SEM of *A. fumigatus* and *C. albicans.*** SEM images of (**a**) *A. fumigatus* and (**b**) *C. albicans*. (**c**) Box and Whisker plot of SEM-measured hyphae width of *A. fumigatus* and cell width and maximum and minimum cell diameters of *C. albicans*. The boxes represent the interquartile range (IQR), with whiskers extending to 1.5 times the IQR. Mean values are represented by white circles, and medians by horizontal lines. (n=38 for *A. fumigatus* and n=14 for both the width and length of *C. albicans* yeast cells.

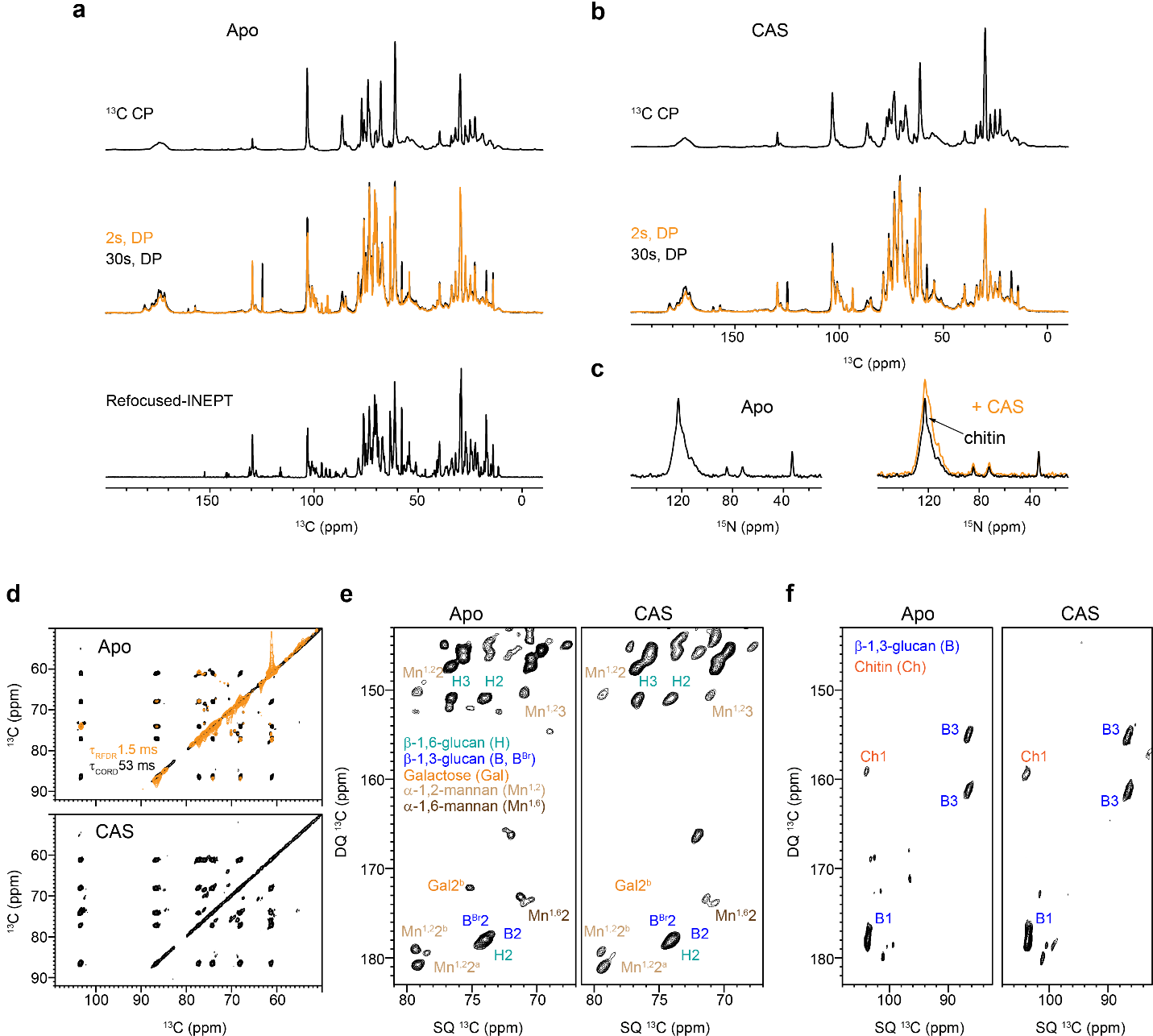

**Supplementary Figure 2. Echinocandin-resistance I.3 strain treated with caspofungin at MIC.** (**a**) 1D ^13^C CP (top), ^13^C DP (middle), and refocused INEPT (bottom) spectra of apo cell wall. (**b**) 1D ^13^C CP (top) and ^13^C DP (bottom) spectra of drug-treated cell wall. (**c**) Comparison of 1D ^15^N CP spectra of cell walls with and without drug. (**d**) Comparison of 2D ^13^C-^13^C 53 ms CORD spectra of cell walls without (top) and with drug (bottom). 2D ^13^C-^13^C 1.5 ms RFDR spectrum overlay on CORD spectrum for apo cell wall. (**e**) 2D ^13^C-^13^C refocused DP-J INADEQUATE spectra of AR387 strain with and without drug cell walls. (**f**) 2D ^13^C-^13^C CP INADEQUATE spectra of AR387 strain with and without drug detected rigid cell wall polysaccharides.

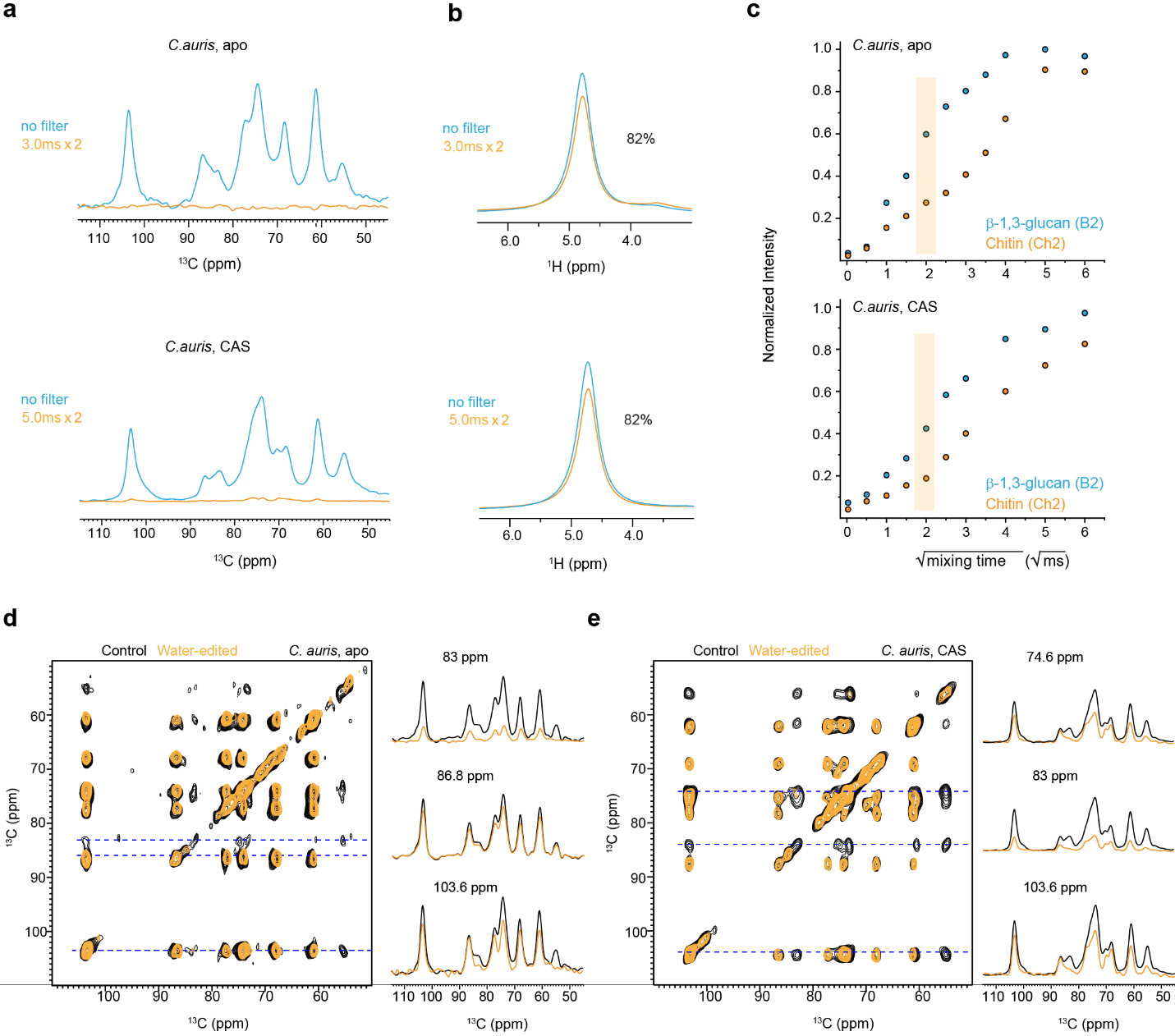

**Supplementary Figure 3. Water-edited spectra of *C. auris* to access polymer hydration.** (**a**) ^1^H-T_2_ filtered (orange) and control (blue) ^13^C spectra are shown for apo (top) and caspofungin-treated (bottom) *C. auris* samples. No spin diffusion was applied. Approximately 92% of carbohydrate ^13^C signals were removed by the ^1^H-T_2_ filter. (**b**) ^1^H-T_2_ filtered (orange) and control (blue) ^1^H NMR spectra, with 82% of water signal retained for both samples after the ^1^H-T_2_ filter. (**c**) Representative water-to-polysaccharide ^1^H spin diffusion buildup curves. Overlay of 2D water-edited (orange) and control (black) ^13^C-^13^C correlation spectra of (**d**) apo sample and (**e**) caspofungin-treated *C. auris*. Representative 1D slices extracted from the 2D ^13^C-^13^C correlation spectra are shown for each sample. The control data are displayed as black solid lines, and the water-edited spectra are plotted in orange. All spectra were measured on a 400 MHz spectrometer at 10 kHz MAS.

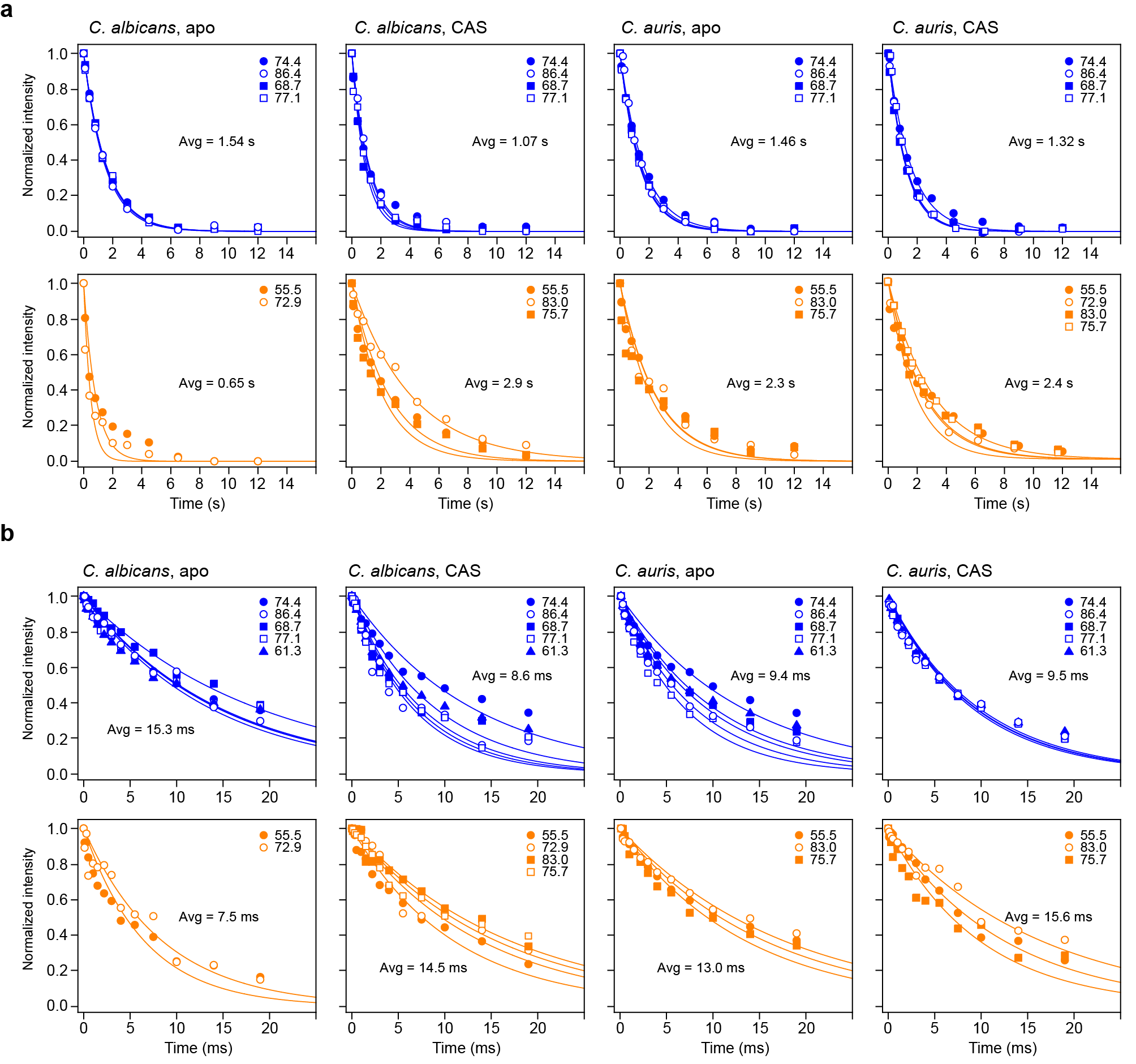

**Supplementary Figure 4.** **Relaxation curves of polysaccharides in *Candida* cell walls.** The relaxation decay curves are shown separately for (**a**) ^13^C T_1_ and (**b**) ^1^H T_1ρ_ of the *Candida* species (SC5314 and AR386) with and without drug-treated cell walls.

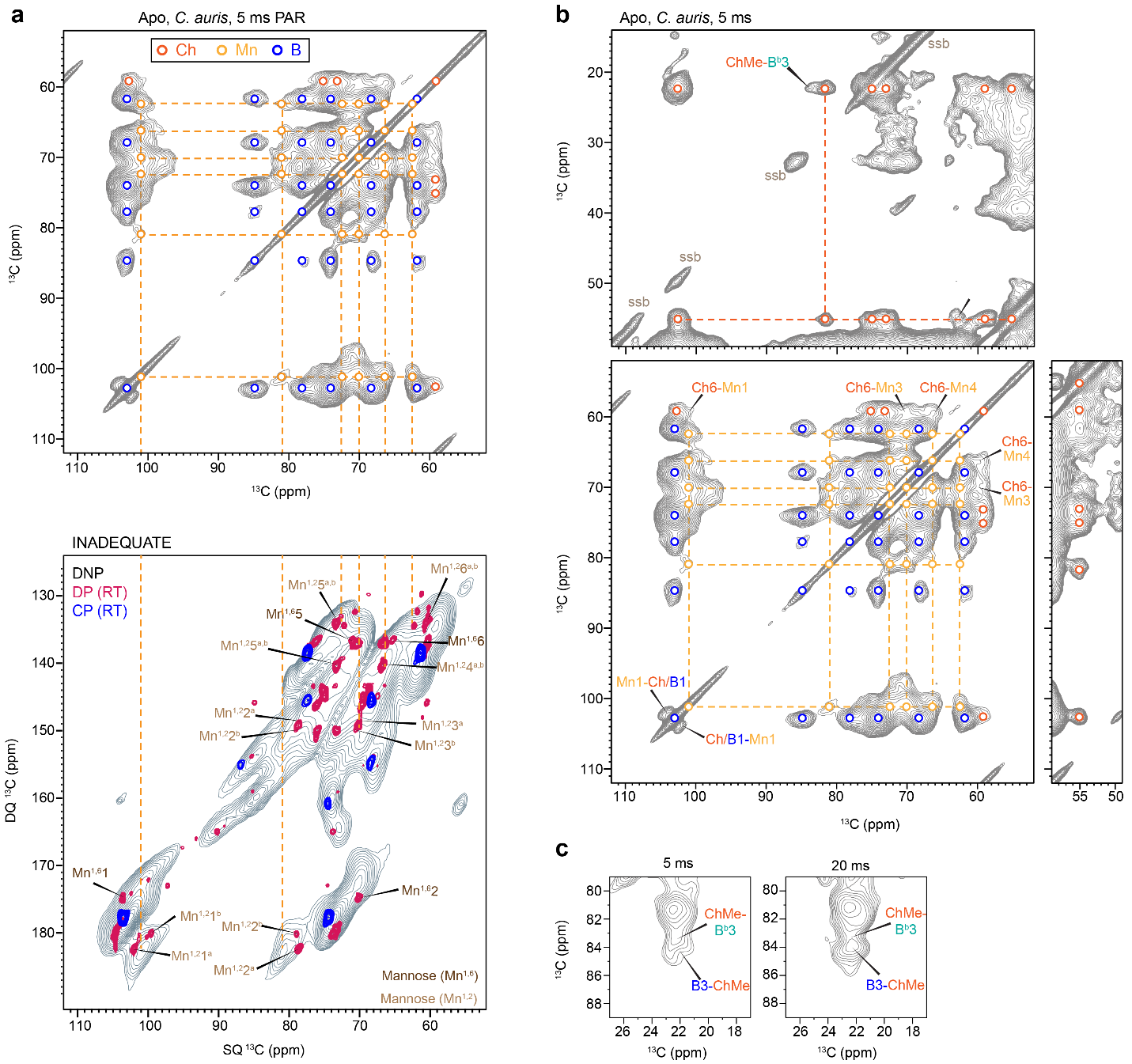

**Supplementary Figure 5. Intermolecular interactions of apo *C. auris* cell walls.** (**a**) DNP 2D ^13^C correlation spectra measured with 5 ms PAR (top) pulse sequences on apo *C. auri*s AR386. Intramolecular cross peaks within each molecule are shown using open circles for chitin (orange), β-1,3-glucan (blue) and mannan (orange). For comparison, refocused J-INADEQUATE spectra of *C. albicans* measured under DNP condition (grey) is shown as the bottom panel, with comparison to room-temperature spectra of *C. albicans* measured with CP (blue) and DP (magenta). Signals of α-1,2-linked mannose residues aligned well with the third-type of signals identified in 5 ms PAR in addition to chitin and β-1,3-glucan. (**b**) Intermolecular cross peaks identified in DNP 5 ms PAR spectrum of *C. auris*. (**c**) Zoomed-in region of 5 ms and 20 ms PAR spectra of *C. auris* showing interactions between chitin methyl and β-1,3-glucan carbon 3.

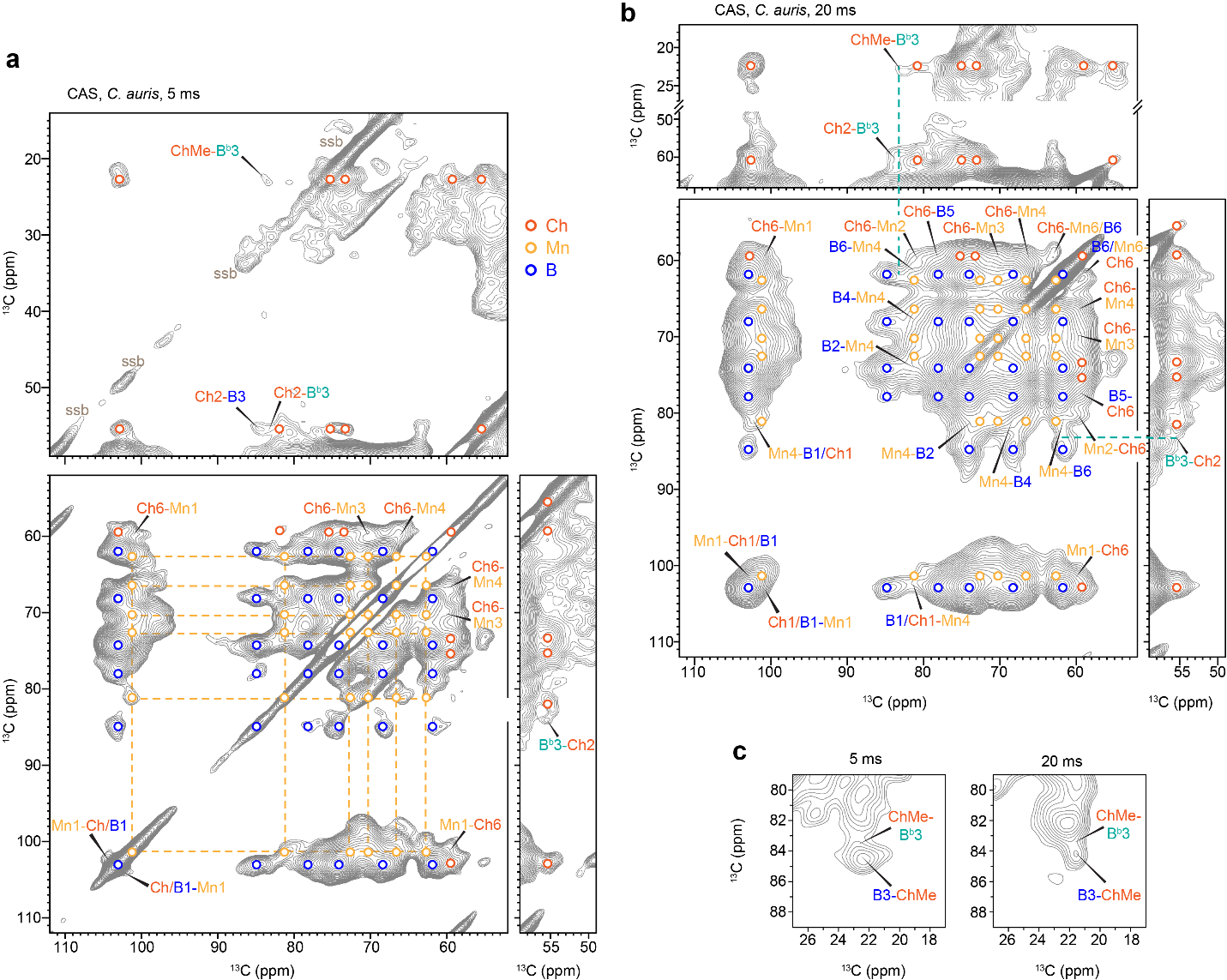

**Supplementary Figure 6. Intermolecular interactions of caspofungin-treated *C. auris* cell walls.** (**a**) DNP-enhanced 5 ms PAR spectrum of caspofungin-treated *C. auris* sample. Intramolecular cross peaks within each molecule are shown using open circles for chitin (orange), β-1,3-glucan (blue), and mannan (orange). (**b**) Intermolecular cross peaks identified in DNP 5 ms PAR spectrum. (**c**) Zoomed-in region of 5 ms and 20 ms PAR spectra showing interactions between chitin methyl and β-1,3-glucan carbon 3.

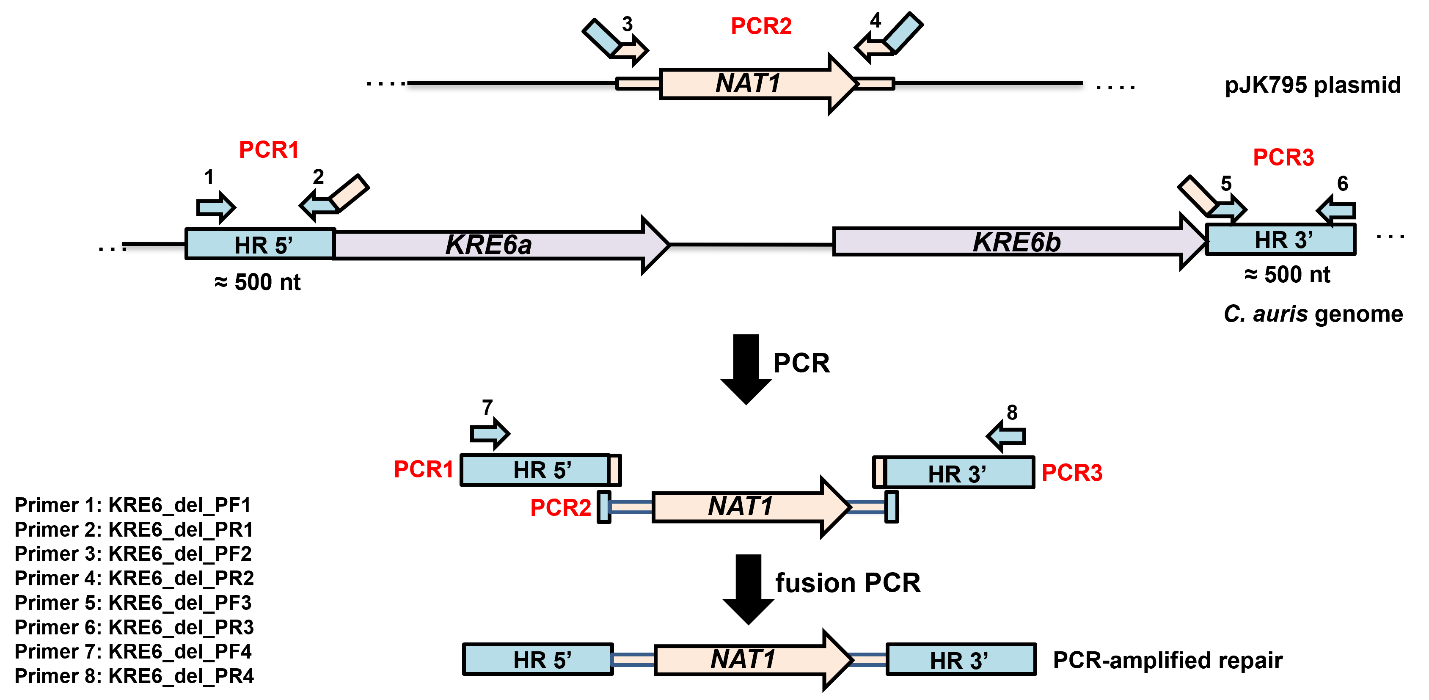

**Supplementary Figure 7. Construction of the *C. auris* *kre6ab*∆ strain.** Schematic view of designed fusion PCRs to obtain a PCR-amplified repair fragment containing *NAT1* and the two-sided homolog regions (HR) with about 500 bp in the upstream region of *KRE6a* and downstream region of *KRE6b*. The fusion PCRs were carried out with overlapping primers, as indicated. Primer 2 is reverse complemented to primer 3, and primer 4 is reverse complemented to primer 5.

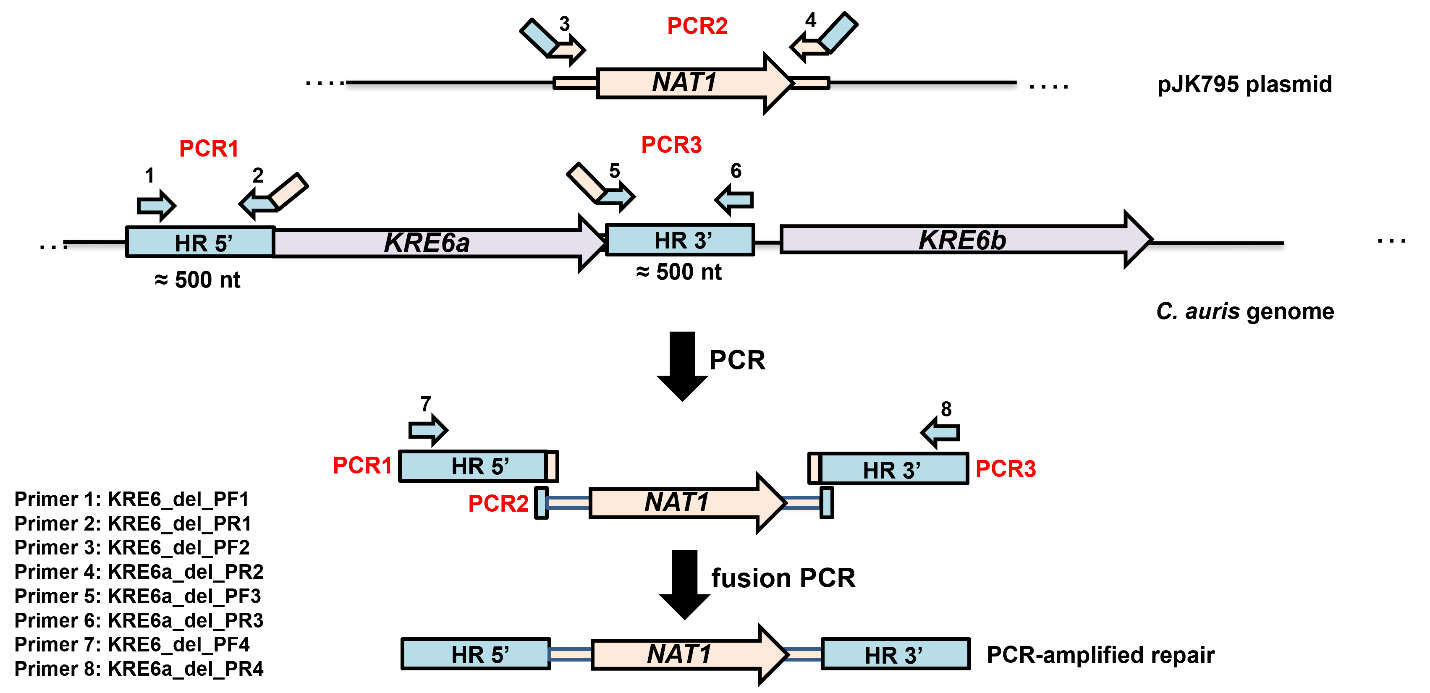

**Supplementary Figure 8. Construction of the *C. auris* *kre6a*∆ strain.** Schematic view of designed fusion PCRs to obtain a PCR-amplified repair fragment to delete *KRE6a*. The fragment contains *NAT1* and the two-sided homolog regions (HR) with about 500 bp in the upstream and downstream regions of *KRE6a*. The fusion PCRs were carried out with overlapping primers, as indicated. Primer 2 is reverse complemented to primer 3, and primer 4 is reverse complemented to primer 5.

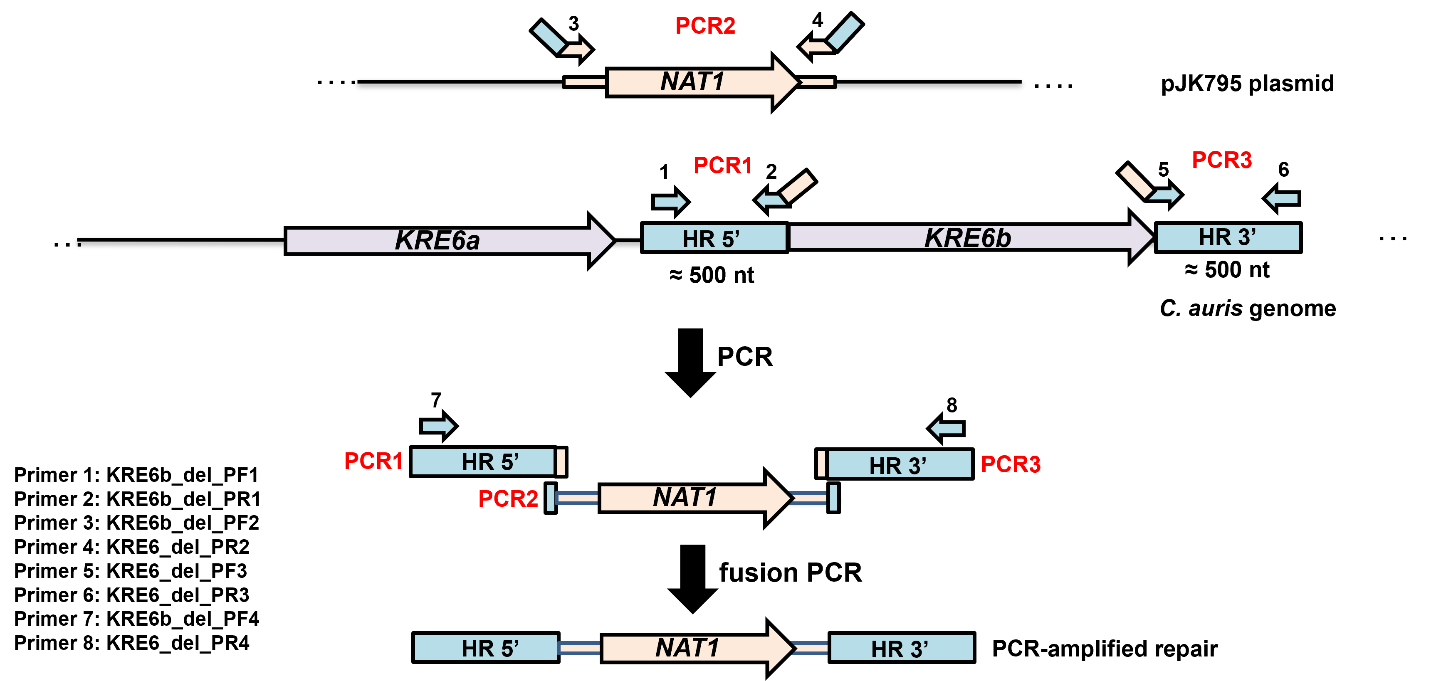

**Supplementary Figure 9. Construction of the *C. auris* *kre6b*∆ strain.** Schematic view of designed fusion PCRs to obtain a PCR-amplified repair fragment to delete *KRE6b*. The fragment contains *NAT1* and the two-sided homolog regions (HR) with about 500 bp in the upstream and downstream regions of *KRE6b*. The fusion PCRs were carried out with overlapping primers, as indicated. Primer 2 is reverse complemented to primer 3, and primer 4 is reverse complemented to primer 5.

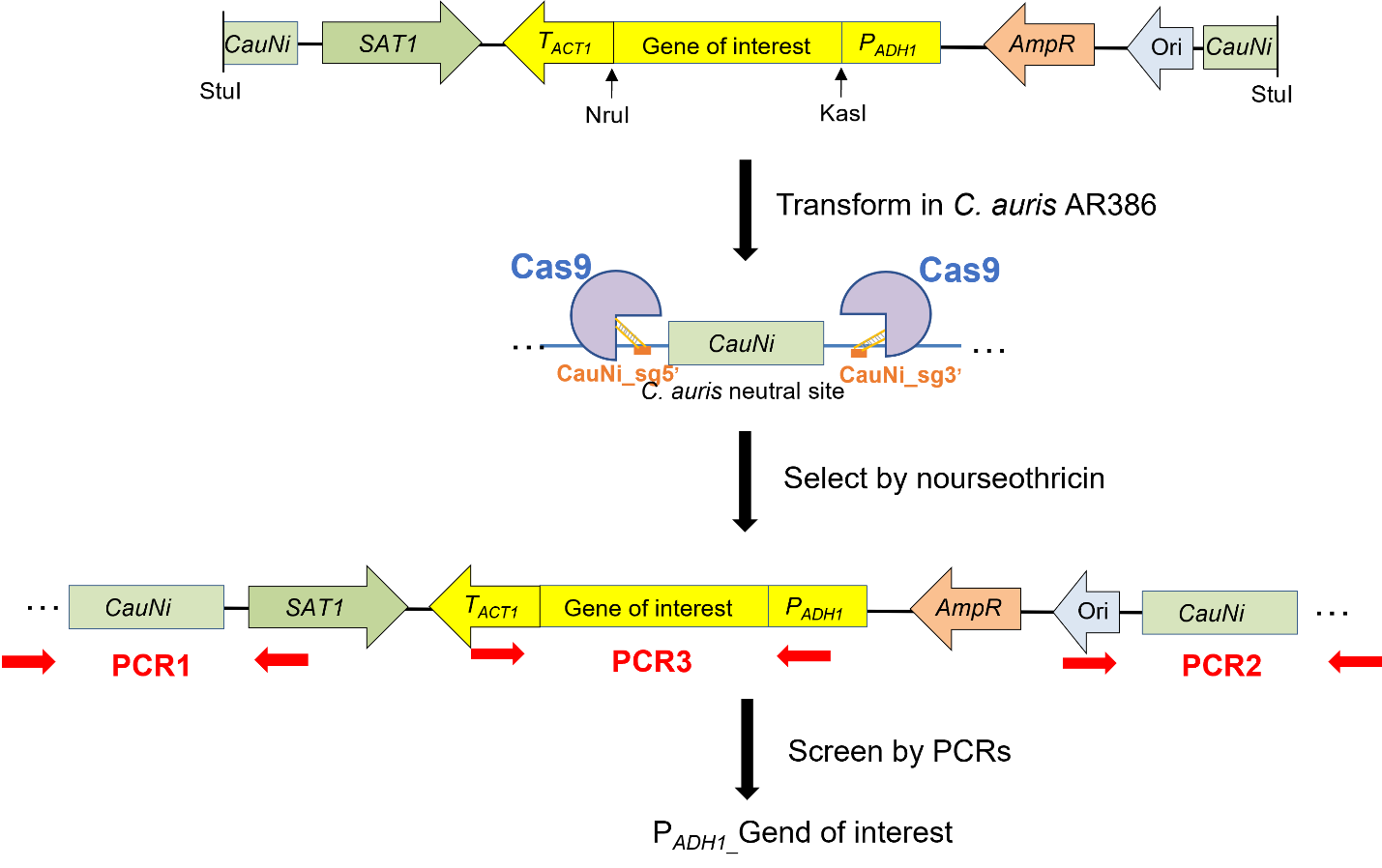

**Supplementary Figure 10. Construction of the *C. auris* P*_ADH1_*_*KRE6a* and P*_ADH1_*_*KRE6b* strains.** Schematic view of the overexpression system in *C. auris*. The plasmid contains the promotor P*_ADH1_*, the gene of interest (*KRE6a* or *KRE6b*), the terminator of T*_ACT1_*, the *SAT1* cassette (nourseothricin resistance) and the *C. auris* neutral site *CauNi*. The restriction sites KasI and NruI were used to insert the nucleotide sequences of gene of interest. The plasmid was linearized by StuI and transformed via electroporation in the wild-type AR386 strain with CRISPR-Cas9 method, by targeting the upstream and the downstream regions of *CauNi* with the nucleotide-specific guide RNAs CauNi_sg5’ and CauNi_sg3’. The transformants were screened by performing three PCRs to verify the proper integration of the plasmid in the genome of *C. auris*.

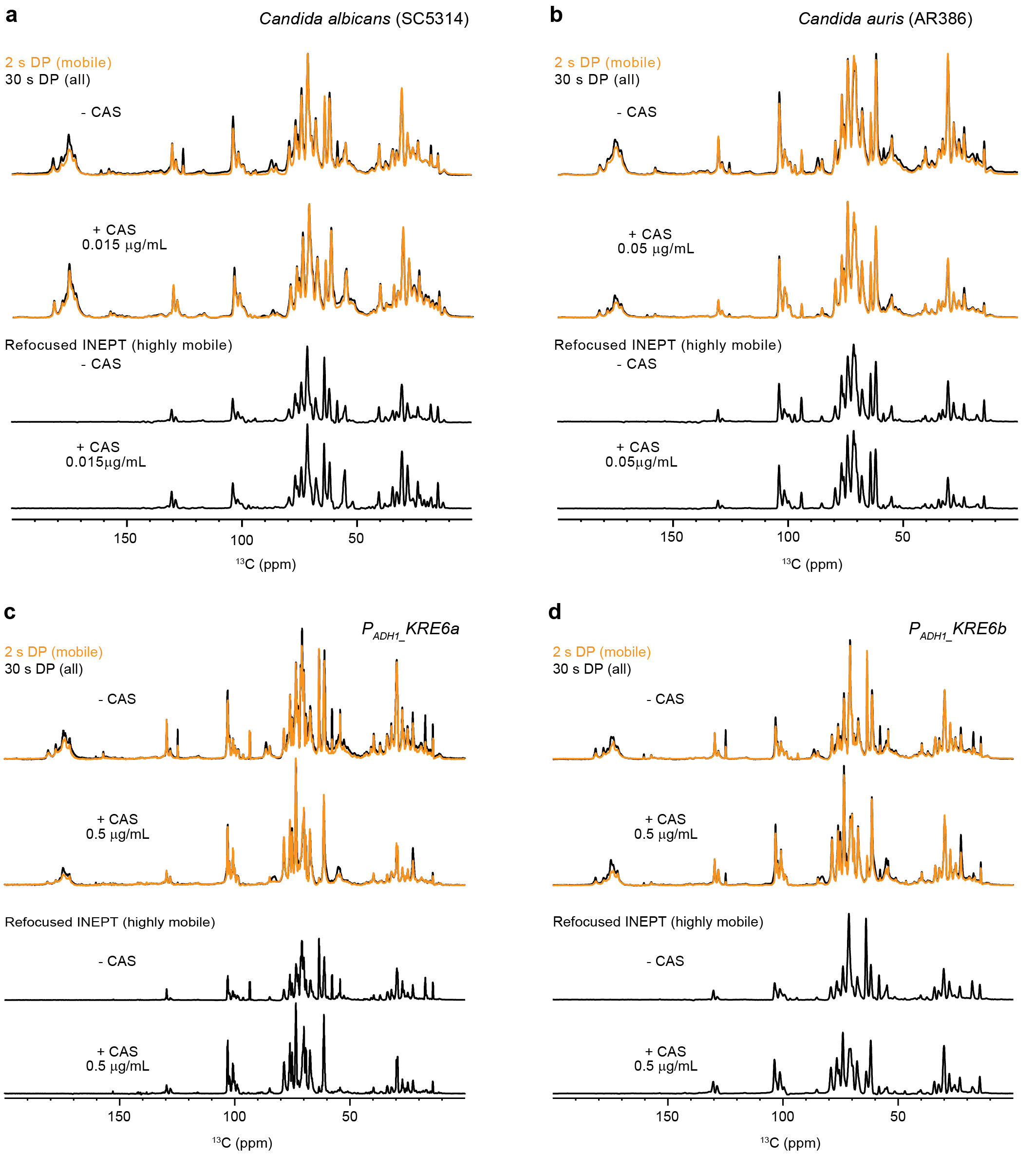

**Supplementary Figure 11. Protein and lipid content in the mobile phase of *C. auris* cell walls.** 1D ^13^C spectra of (**a**) *C. albicans* and (**b**) *C. auris* with and without CAS treatment. The spectra include 1D ^13^C 2 s DP for detecting mobile molecules, 30 s DP for quantitatively detecting all molecules, and 1D ^13^C refocused INEPT for detecting highly mobile molecules in *C. albicans* and *C. auris*. The same set of 1D ^13^C spectra were also measured on (**c**) P_ADH1__*KRE6a* and (**d**) P_ADH1__*KRE6b* with and without CAS treatment. The spectra include 1D ^13^C 2 s DP for detecting mobile molecules, 30 s DP for quantitatively detecting all molecules, and 1D ^13^C refocused INEPT for detecting highly mobile molecules in *C. auris* *KRE6* mutants.

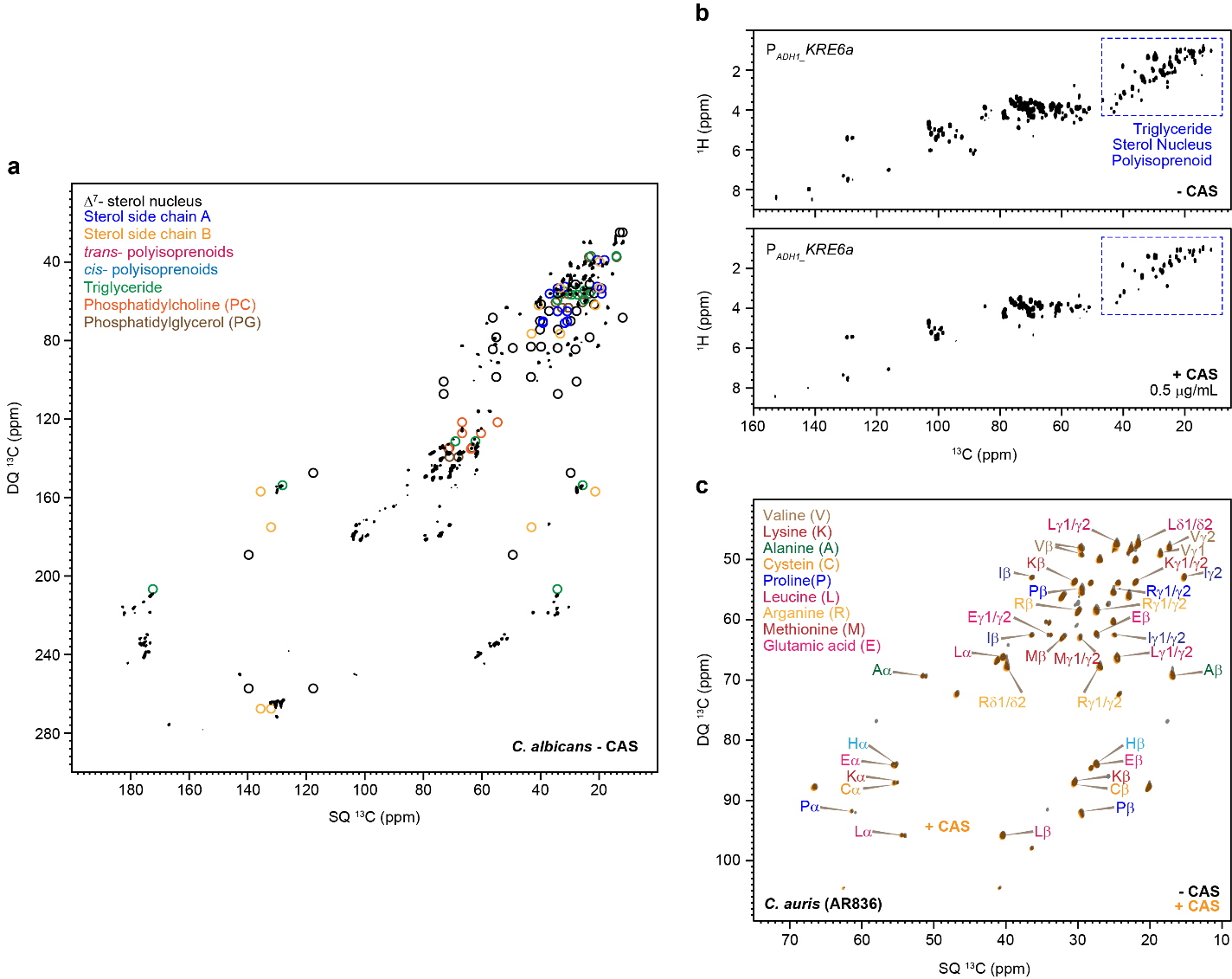

**Supplementary Figure 12. Protein and lipids in the mobile fraction of *Candida* cell walls.** (**a**) Overlay of 2D ^13^C-^13^C refocused DP J-INADEQUATE spectra of *C. albicans* with drug sample (black) with simulated spectra. The simulated spectra were plotted using the chemical shifts reported in recent NMR studies of lipid components (sterol, polyisoprenoid, and triglycerides) and model phospholipids POPC and POPG.^1-5^ (**b**) 2D ^1^H-^13^C refocused INEPT spectra of *C. auris* P_ADH1__*KRE6a* with and without drug show the highly mobile protein, lipids, and polysaccharides. Protein and lipids are shown in blue color dashed boxes. (**c**) 2D ^1^H-^15^N HETCOR spectra showing amide and amine signals. (**c**) 2D ^13^C-^13^C refocused DP J-INADEQUATE spectra of *C. auris* AR386 showing signals of mobile proteins, with highly overlapping signals that indicate similar protein structures^6^.

**Supplementary Table 1. The average cell wall thickness of comparable-sized yeast cells.** Results are described as the average and standard deviation of 100 measurements for each sample. A source data file is provided to document each single reading.

| Sample | Strain | Average cell wall thickness (nm) | |
| --- | --- | --- | --- |
|  |  | apo | +CAS |
| *C. albicans* | SC5314 | 140±11 | 185±16 |
| *C. auris* | AR836 | 158±11 | 162±13 |
|  | I.3 | 70±17 | 75±17 |
|  | P_ADH1__*KRE6a* | 114±23 | 183±28 |
|  | P_ADH1__*KRE6b* | 116±26 | 175±21 |

**Supplementary Table 2.** **^13^C and ^15^N chemical shifts of polysaccharides in cell walls at room temperature and at DNP conditions.** Branched (Br). Not applicable (/). Unidentified (-). Minor (m). The referencing scale is TMS scale.

| Carbohydrate | | C1 | C2 | C3 | C4 | C5 | C6 | CO | CH_3_ | N | Experimental method | Sample | Reference |
| --- | --- | --- | --- | --- | --- | --- | --- | --- | --- | --- | --- | --- | --- |
| Room-temperature solid-state NMR | | | | | | | | | | | | | |
| β-1,3-glucan |  | 103.7 | 74.5 | 86.8 | 68.3 | 77.5 | 61.2 | / | / | / | 53 ms CORD | *C. albicans* apo | Shim *et al*. 2007 ^7^  Fairweather *et al*. 2009 ^8^  Saito *et al*. 1979 ^9^ |
|  | m | 103.6 | 74.6 | 85.2 | 68.3 | 77.4 | 61.1 | / | / | / |  | *C. albicans*,  *C. auris* |  |
|  | m | 103.3 | 74.4 | 85.3 | 68.3 | 77.5 | 61.1 | / | / | / |  |  |  |
| Chitin | | 103.8 | 55.0 | 73.2 | 83.2 | 75.8 | 60.9 | 174.9 | 22.9 | 122.3 | 53 ms CORD | *C. albicans* | Fernando *et al.* 2021 ^10^ |
|  |  | 101.5 | 55.2 | 72.4 | 84.5 | 75.1 | 60.2 | 175.7 | 23.2 | 122.3 |  | *C. auris* |  |
| β-1,3,6-glucan | Br | 103.1 | 73.2 | 85.5 | 68.7 | 75.7 | 69.2 | / | / | / | Refocused DP J-INADEQUTE | *C. albicans*, *C. auris* | Lowman *et al*. 2011 ^11^ |
| β-1,6-glucan | | 103.9 | 74.2 | 76.3 | 70.6 | 75.4 | 69.6 | / | / | / |  |  |  |
| α-1,2-Mannan | a | 101.4 | 79.1 | 71.0 | 67.5 | 73.8 | 61.9 | / | / | / |  | all samples | Latgé *et al*.1994 ^12^  Chakraborty *et al*. 2021 ^13^ |
|  | b | 99.2 | 79.1 | 71.4 | 67.8 | 74.1 | 61.9 | / | / | / |  |  |  |
| α-1,6-Mannan | | 102.7 | 70.8 | 73.8 | 67.7 | 71.2 | 67.3 | / | / | / |  |  |  |
| Galactose | a | 90.2 | 74.2 | - | - | - | - | / | / | / |  | *C. albicans* apo, CAS | Fontaine *et al*. 2011 ^14^ |
|  | b | 96.8 | 74.9 | 76.8 | 70.6 | 72.3 | 61.5 | / | / | / |  | all samples |  |
|  | d | 94.8 | 72.0 | - | - | - | - | / | / | / |  |  |  |
| Glucose | a | 92.9 | 72.1 | 73.8 | 70.5 | 72.0 | 63.9 | / | / | / |  | all samples |  |
|  | b | 96.8 | 74.9 |  |  |  |  |  |  |  |  |  |  |
| MAS-DNP | | | | | | | | | | | | | |
| β-1,3-glucan |  | 103.0 | 74.1 | 85.0  83.9 | 68.1 | 77.8 | 61.6 |  |  |  | DNP 5 ms PAR | *C. auris,* apo, CAS |  |
| Chitin |  | 102.5 | 55.2 | 72.8 | 81.9 | 74.8 | 59.1 | 175.9  174.9  172.9 | 22.3 |  |  |  |  |
| α-1,2-Mannan |  | 101.4 | 81.1 | 70.0 | 66.3 | 72.3 | 62.5 |  |  |  |  |  |  |
| α-1,2-Mannan (m) |  | 101.9 | 81.9 |  |  |  |  |  |  |  |  |  |  |

**Supplementary Table 3.** **^1^H and ^13^C chemical shifts of *C. albicans* and *C. auris* polysaccharides from ^1^H-detection experiments.** For each carbon site, the ^13^C and ^1^H chemical shifts are shown in the top and bottom rows, respectively. The referencing scale is TMS scale for ^13^C, and DSS for ^1^H. ^13^C and ^1^H sites with ambiguity are underlined: for Gl, the carbon numbering of C2, 3, 4 is uncertain.

| Carbohydrates | forms | C1 | C2 | C3 | C4 | C5 | C6 | Reference |
| --- | --- | --- | --- | --- | --- | --- | --- | --- |
| Rigid molecules | | | | | | | | |
| β -1,3-glucan (B) |  | 103.9  5.0 | 74.44  3.74 | 86.89  3.71 | 68.52  3.71 | 77.6  3.54 | 61.54  3.94 | Chakraborty *et al.* 2021 ^13^ |
| Chitin (Ch) |  | --- | 55.5  4.4 | --- | 83.3  3.8 | 75.8  3.6 | --- | Fernando *et al.* 2021 ^10^ |
| Mobile molecules | | | | | | | | |
| β-1,3-glucan (B) | a | 103.6  4.55 | 74.1  3.3 | 85.2  3.75 | 69.3  4.2 | 76.0  3.52 | 61.6  3.75 | Shim *et al.* 2007 ^7^  Fairweather *et al.* 2009 ^8^  Saito *et al.* 1979 ^9^ |
|  | b | --- | 74.65  3.4 | 85.12  4.12 | 69.9  4.2 | --- | 61.95  3.9 |  |
|  | c | --- | 74.49  3.52 | 86.74  4.28 | 71.59  4.43 | --- | 62.5  3.8 |  |
| β-1,3,6-glucan (Br) |  | --- | --- | 84.7  4.3 | --- | 75.44  3.6 | 69.6  3.8 | Lowman *et al.* 2011 ^11^ |
| β-1,6-glucan (H) |  | 103.54  4.52 | 74.15  3.3 | 76.3  3.5 | 70.4  3.89 | --- | 69.9  3.87 |  |
| α-1,6-Mannan (Mn^1,6^) |  | 102.7  5.14 | 70.8  3.4/4.0 | 73.8  3.34/3.7 | 67.7  3.6 | 70.8  3.8 | --- | Latge *et al.* 1994 ^12^  Chakraborty *et al.* 2021 ^13^ |
| α-1,2-Mannan (Mn^1,2^) | a | 101.4  5.28 | 79.19  4.11 | 70.89  3.94 | 67.8  3.7 | 73.98  3.74 | 62.1  3.9 | Kuraoka *et al.* 2021 ^15^ |
|  | b | 98.97  5.08 | 79.51  4.0 | 71.19  3.95 | 67  3.71 | 73.4  3.68 | 61.98  3.94 |  |
|  | c | 100.74  5.16 | 78.22  4.28 | 70.12  3.94/4.19 | 68.09  3.6 | 73.98  3.75 | 61.73  3.77 |  |
|  | d | 101.45  5.36 | 79.12  4.11 | 70.66  3.92 | 67.88  3.6 | 73.61  3.64 | 62.0  3.67 | Kuraoka *et al.* 2018 ^16^ |
|  | e | 102.7  5.04 | 79.13  3.91/4.1 | 71.2  3.8 | 67.7  3.6 | 74.23  3.34 | 61.9  3.7, 3.9 | Kuraoka *et al.* 2021^15^ |
| Galactose/Glucose or their derivatives  (Gl) | a | 90.04  5.9 | 85.4  4.1/3.7 | 74.5  4.34 | 70.27  4.2 | --- | 61.84  3.8, 3.9 | Fontaine *et al.* 2011 ^14^ |
|  | b | 97.06  5.45 | 71.08  4.02 | 74.36  3.2 | 67.03  3.72 | --- | 61.82  3.7, 3.9 | Archbald et al. 1981^17^ |
|  | c | 94.66  4.9 | 72.5  3.8 | 73.59  3.65 | 67.42  3.58 | --- | 61.9  3.7, 3.8 | Archbald et al. 1981 ^17^  Fontaine *et al.* 2011 ^14^ |
|  | d | 94.98  5.18 | 72.5  3.81 | 71.3  3.85 | 67.6  3.66 | --- | 61.9  3.7.3.8 | Archbald et al. 1981 ^17^  Fontaine *et al.* 2011^14^ |
|  | e | 89.24  6.05 | 86.74  4.2 | 74.8  4.34/3.4 | 71.39  4.4 | --- | 62.59  3.8, 3.9 | Fontaine *et al.* 2011 ^14^ |
| Glucose  (Glc) | a (α) | 93.07  5.237 | 72.4  3.5 | 72.4  3.5 | 70.4  3.4 | --- | 61.56  3.84, 3.76 | Roslund et al. 2008  Archbald *et al.* 1981 ^17^ |
|  | b (β) | 96.84  4.65 | 74.9  3.25 | 76.58  3.47 | 70.48  3.4 | --- | 61.79  3.9,3.73 |  |

**Supplementary Table 4. The molar composition of rigid polysaccharides.** The numbers are estimated using integrals (volume) of cross peaks in 2D ^13^C-^13^C 53 ms CORD spectra. The average integrals of cross-peaks of each polysaccharide are shown. Error bars are standard errors.

| Sample | | Polysaccharide | | | | | |
| --- | --- | --- | --- | --- | --- | --- | --- |
| Strain | Condition | β-1,3-glucan | | | Chitin | β-1,6-glucan | Mannan |
|  |  | a | b | c |  |  |  |
| *C. albicans* | apo | 87±11 | 1.1±0.2 | 6±1 | 5±2 | 1.1±0.1 | ND |
| (SC5314) | +CAS | 46±6 | 0.4±0.2 | ND | 44±8 | 2.6±0.2 | 7±2 |
| *C. auris* | apo | 63±6 | 6±2 | 11±3 | 18±4 | 1.3±0.1 | 1.1±0.2 |
| (AR386) | +CAS | 37±5 | 1.0±0.3 | 3.0±0.8 | 48±7 | 5.0±0.4 | 5.5±0.8 |
| *C. auris* | apo | 77±9 | ND | 3±2 | 5±2 | 9.1±0.7 | 6±2 |
| (I.3) | +CAS | 66±9 | ND | 5±2 | 12±2 | 5.1±0.3 | 12±2 |
| *C. auris* | apo | 85±11 | ND | 2±1 | 8±2 | 5.2±0.8 | ND |
| (P_ADH1__*KRE6a*) | +CAS | ND | ND | ND | 80±13 | 7.1±0.9 | 13±2 |
| *C. auris* | apo | 84±11 | ND | ND | 12±2 | 4.1±0.5 | ND |
| (P_ADH1__*KRE6b*) | +CAS | ND | ND | ND | 82±16 | 10±1 | 8±1 |
| *C. auris* | apo | 95±13 | ND | ND | ND | 5±2 | ND |
| (*kre6a∆*) | +CAS | 12±2 | ND | ND | 79±13 | 4±1 | 5.0±0.8 |
| *C. auris* | apo | 99.3±9.7 | ND | ND | ND | 0.7±0.3 | ND |
| (*kre6b∆*) | +CAS | 38±6 | ND | ND | 49±10 | 7±2 | 6±1 |

The area of the following well-resolved cross peaks 53 ms CORD spectra are used:

β-1,3 (a): the average of C1-C2/3/4/5 and C3-C2/4/5/6.

β-1,3 (b): the average of C1-C5, C2-5, C4-5, and C6-5.

β-1,3 (c): the average of C3-C2/4/5/6.

β-1,6: the average of C3/5-C4 and C5-C6.

Chitin: the average of C1-2/4/5, C3-C2, C4-C2/3/5, C5-C2, and C6-C2.

Mannan: the average of C2-C5 and C2-C3.

**Supplementary Table 5. The molar composition of mobile polysaccharides.** The numbers are estimated using integrals (volume) of cross peaks in 2D ^13^C-^13^C refocused DP-J INADEQUATE spectra. The average integrals of cross-peaks of each polysaccharide are shown. Error bars are standard errors of the peak integrals.

| Sample | | Polysaccharide | | | | | | | | | |
| --- | --- | --- | --- | --- | --- | --- | --- | --- | --- | --- | --- |
| Strain | Condition | β-glucan | | | Mannan^1,2^ | | Mannan^1,6^ | Galactose | | | |
|  |  | β-1,6-glucan | β-1,3-glucan | β-1,3,6-glc | a | b |  | a | b | c | d |
| *C. albicans* | apo | 30±6 | 33±7 | 1.2±0.3 | 14±2 | 5±1 | 8.9±0.8 | 4.0±0.3 | 2.0±0.3 | 1.1±0.5 | 1.1±0.6 |
| (SC5314) | +CAS | 19±3 | 15±4 | 2.6±0.3 | 24±3 | 5±1 | 21±2 | 7.8±0.5 | 1.7±0.2 | 0.9±0.2 | 2.3±0.5 |
| *C. auris* | apo | 33±4 | 24±2 | 1.5±0.3 | 9±1 | 2.9±0.2 | 6±2 | 4.1±0.3 | 7±1 | 4±1 | 8±2 |
| (AR386) | +CAS | 35±3 | 20±1 | 0.7±0.2 | 14±1 | 2.8±0.1 | 9.1±0.9 | 2.2±0.2 | 7±1 | 4±1 | 6±2 |
| *C. auris* | apo | 33±4 | 31±5 | 2.5±0.2 | 11±2 | 6.3±0.6 | 5±2 | 1.2±0.2 | 4.0±0.5 | 5.3±0.5 | 1.0±0.7 |
| (I.3) | +CAS | 30±2 | 22±4 | 3.1±0.3 | 15±2 | 6±1 | 6±2 | 0.65±0.04 | 2.9±0.6 | 0.7±0.7 | 13±1 |
| *C. auris* | apo | 32±2 | 26±5 | 4.6±0.3 | 12±1 | 4.4±0.7 | 7±1 | 1.24±0.06 | 7.3±0.9 | 4.7±0.7 | 1.4±0.3 |
| (P_ADH1__*KRE6a*) | +CAS | 36±7 | 19±6 | 2.8±0.3 | 24±3 | 5.5±0.5 | 11±3 | 0.9±0.1 | 0.36±0.07 | ND | 0.7±0.2 |
| *C. auris* | apo | 32±4 | 16±4 | 6.3±0.4 | 21±2 | 8.7±0.8 | 13.8±0.9 | 1.32±0.07 | 0.51±0.03 | 0.45±0.08 | 0.9±0.1 |
| (P_ADH1__*KRE6b*) | +CAS | 35±6 | 21±5 | 2.5±0.2 | 23±3 | 5.4±0.7 | 11±1 | 1.2±0.2 | ND | 0.12±0.03 | ND |

| *C. auris* | apo | 19±3 | 44±7 | 5.0±0.8 | 7±1 | 4.0±0.5 | 8±1 | ND | 9.6±3.2 | ND | 3.4±0.4 |
| --- | --- | --- | --- | --- | --- | --- | --- | --- | --- | --- | --- |
| (*kre6a∆*) | +CAS | 34±7 | 36±12 | 4±0.6 | 11±3 | 5.0±0.9 | 10±3 | ND | ND | ND | ND |

| *C. auris* | apo | 23±5 | 40±14 | 7±1 | 5±1 | 3.0±0.8 | 6±2 | ND | 11±3 | ND | 5±1 |
| --- | --- | --- | --- | --- | --- | --- | --- | --- | --- | --- | --- |
| (*kre6b∆*) | +CAS | 53±16 | 24±8 | 4.0±0.9 | 7.0±1.2 | 4±1 | 8±2 | ND | ND | ND | ND |

The area of the following well-resolved cross peaks refocused DP-J INADEQUATE spectra are used:

β-1,6: the average of C3, C4, C5, and C6.

β-1,3; the average of C1, C2, C5, and C6

β-1,3, 6: the average of C2, C3, and C4.

Mannan^1,2^ (a and b): the average of C1 and C2.

Mannan^1,6^: the average of C1 and C2.

Galactose (a, b, c, and d): the average of C1 and C2.

**Supplementary Table 6. Water-edited intensities of polysaccharide carbon sites.** The intensity ratios are obtained by comparing the peak intensities in water-edited and control 2D spectra. The average values for each molecule in each sample are highlighted in bold. Error bars are standard deviations propagated from NMR signal-to-noise ratios.

| Polysaccharide | Cross-peak | *C. albicans* (SC5314) | | *C. auris* (AR386) | |
| --- | --- | --- | --- | --- | --- |
|  |  | apo | +CAS | apo | +CAS |
| β-1,3-glucan | B1-3 | 0.8±0.1 | - | 0.6±0.1 | 0.28±0.06 |
|  | B1-5 | 0.7±0.1 | 0.8±0.3 | 0.5±0.1 | 0.25±0.05 |
|  | B1-2 | 0.73±0.08 | 0.6±0.1 | 0.54±0.07 | 0.19±0.02 |
|  | B1-4 | 0.7±0.1 | 0.7±0.2 | 0.6±0.1 | 0.29±0.05 |
|  | B1-6 | 0.7±0.1 | 0.5±0.1 | 0.7±0.1 | 0.17±0.04 |
|  | B3-1 | 0.7±0.1 | - | 0.7±0.2 | 0.26±0.04 |
|  | B3-5 | 0.7±0.2 | 0.8±0.3 | 0.6±0.2 | 0.26±0.06 |
|  | B3-2 | 0.8±0.1 | 0.6±0.2 | 0.5±0.1 | 0.22±0.05 |
|  | B3-4 | 0.7±0.1 | 0.9±0.3 | 0.5±0.1 | 0.24±0.06 |
|  | B3-6 | 0.8±0.2 | 0.8±0.4 | 0.5±0.2 | 0.29±0.07 |
|  | B5-1 | 0.8±0.1 | 0.8±0.2 | 0.6±0.1 | 0.23±0.04 |
|  | B5-3 | 0.8±0.2 | 0.8±0.3 | 0.6±0.2 | 0.22±0.05 |
|  | B5-2 | 0.7±0.1 | 0.5±0.1 | 0.6±0.1 | 0.15±0.04 |
|  | B5-4 | 0.7±0.1 | 0.8±0.2 | 0.6±0.1 | 0.25±0.03 |
|  | B5-6 | 0.67±0.07 | 0.7±0.1 | 0.56±0.08 | 0.21±0.03 |
|  | B2-1 | 0.66±0.07 | 0.61±0.09 | 0.58±0.06 | 0.20±0.02 |
|  | B2-3 | 0.8±0.2 | - | 0.7±0.1 | 0.23±0.05 |
|  | B2-5 | 0.9±0.2 | 0.7±0.2 | 0.6±0.1 | 0.19±0.04 |
|  | B2-4 | 0.7±0.1 | 0.7±0.2 | 0.60±0.09 | 0.23±0.04 |
|  | B2-6 | 0.6±0.1 | 0.48±0.09 | 0.5±0.1 | 0.16±0.03 |
|  | B4-1 | 0.64±0.09 | 0.7±0.2 | 0.51±0.09 | 0.24±0.04 |
|  | B4-3 | 0.5±0.1 | 0.7±0.2 | 0.6±0.1 | 0.25±0.05 |
|  | B4-5 | 0.7±0.1 | 0.8±0.2 | 0.6±0.1 | 0.27±0.04 |
|  | B4-2 | 0.61±0.08 | 0.6±0.1 | 0.50±0.09 | 0.19±0.03 |
|  | B4-6 | 0.71±0.08 | 0.6±0.1 | 0.50±0.09 | 0.26±0.04 |
|  | Average | 0.71 | 0.68 | 0.57 | 0.23 |
| Chitin | Ch1-4 | 0.5±0.2 | 0.3±0.1 | 0.6±0.2 | 0.06±0.04 |
|  | Ch1-5 | - | 0.6±0.1 | 0.36±0.06 | 0.17±0.02 |
|  | Ch1-2 | - | 0.17±0.07 | 0.3±0.2 | 0.08±0.03 |
|  | Ch4-1 | - | 0.2±0.1 | - | 0.07±0.04 |
|  | Ch4-5 | 0.7±0.4 | 0.3±0.2 | - | 0.08±0.02 |
|  | Ch4-2 | - | - | - | 0.08±0.04 |
|  | Ch5-4 | 0.3±0.1 | 0.28±0.07 | 0.3±0.1 | 0.04±0.02 |
|  | Ch5-2 | - | 0.22±0.09 | - | 0.05±0.03 |
|  | Ch3-4 | 0.6±0.3 | 0.25±0.09 | 0.2±0.1 | 0.08±0.03 |
|  | Ch3-2 | - | 0.3±0.1 | 0.2±0.1 | 0.07±0.02 |
|  | Ch2-1 | - | 0.16±0.06 | 0.1±0.1 | 0.10±0.04 |
|  | Ch2-4 | - | 0.19±0.09 | 0.3±0.4 | 0.06±0.04 |
|  | Ch2-5 | - | 0.25±0.09 | 0.1±0.2 | 0.04±0.03 |
|  | Ch2-3 | - | 0.25±0.05 | 0.2±0.2 | 0.09±0.03 |
|  | Ch2-1 | - | 0.2±0.1 | - | 0.09±0.07 |
|  | Average | 0.53 | 0.27 | 0.27 | 0.08 |

**Supplementary Table 7. ^1^H-T_1ρ_ and ^13^C-T_1_ relaxation times of polysaccharides in cell walls.** Data are shown for the *C. albicans* and *C. auris* samples, with and without treatment by caspofungin. The average values for each molecule in each sample are highlighted in bold. The data were measured using 1D ^13^C relaxation experiments. The data are fit using single exponential equations: $I\left( t \right)=e^{\frac{-t}{T_{1}}}$. Error bars are standard deviations of the fit parameters.

| Polysaccharide | Chemical shift | *C. albicans* (SC5314) | | | | *C. auris* (AR386) | | | |
| --- | --- | --- | --- | --- | --- | --- | --- | --- | --- |
|  |  | apo | | +CAS | | apo | | +CAS | |
|  |  | ^1^H-T_1ρ_ (ms) | ^13^C-T_1_ (s) | ^1^H-T_1ρ_ (ms) | ^13^C-T_1_ (s) | ^1^H-T_1ρ_ (ms) | ^13^C-T_1_ (s) | ^1^H-T_1ρ_ (ms) | ^13^C-T_1_ (s) |
| β-1,3-glucan | 86.4 | 14.5±0.6 | 1.47±0.03 | 6.6±0.6 | 1.19±0.06 | 9.4±0.6 | 1.36±0.03 | 8.1±0.5 | 1.25±0.02 |
|  | 77.1 | 14.5±0.9 | 1.54±0.04 | 7.0±0.5 | 1.03±0.07 | 9.1±0.5 | 1.40±0.03 | 7.0±0.6 | 1.25±0.02 |
|  | 74.4 | 14.8±0.7 | 1.58±0.02 | 13.1±0.9 | 1.14±0.09 | 9.5±0.5 | 1.63±0.05 | 13±1 | 1.57±0.07 |
|  | 68.7 | 18.8±0.5 | 1.56±0.03 | 7.5±0.6 | 0.92±0.05 | 9.4±0.4 | 1.47±0.03 | 9.4±0.6 | 1.20±0.04 |
|  | 61.3 | 13.7±0.9 | - | 9.0±0.7 | **-** | 9.8±0.5 | **-** | 10.2±0.8 | **-** |
|  | Average | 15.3 | 1.5 | 8.6 | 1.1 | 9.4 | 1.5 | 9.5 | 1.3 |
| Chitin | 83 | - | - | 17.0±0.9 | 4.1±0.2 | 16.5±0.9 | 2.4±0.3 | 17.7±0.9 | 3.0±0.2 |
|  | 75.7 | - | - | 16±1 | 2.1±0.2 | 9.9±0.9 | 2.1±0.3 | 13.5±0.8 | 2.4±0.2 |
|  | 72.9 | 8.5±0.8 | 0.5±0.1 | 14±1 | - | - | - | - | 1.8±0.2 |
|  | 55.5 | 6.5±0.6 | 0.8±0.1 | 11.0±0.7 | 2.6±0.2 | 12.5±0.4 | 2.5±0.2 | 15.6±0.8 | 2.5±0.2 |
|  | Average | 7.5 | 0.65 | 14.5 | 2.9 | 13.0 | 2.3 | 15.6 | 2.4 |

**Supplementary Table 8. Primers and RNAs guides used in this study.**

| **Primer / guide RNA** | **Aim** | **Sequence (5’ 🡪 3’)** |
| --- | --- | --- |
| ***KRE6a*/*b* deletion** | | |
| KRE6_del_PF1 | *KRE6a*/*b* and *KRE6a* deletion cassette construction, *KRE6a* deletion verification | TAT GCA GTA CGC GTG AAA ACT GC |
| KRE6_del_PR1 | *KRE6a*/*b* and *KRE6a* deletion cassette construction | GTA TTC TGG GCC TCC ATG TCA GCG TTT GGG GAT GAA GAT GG |
| KRE6_del_PF2 | *KRE6a*/*b* and *KRE6a* deletion cassette construction | CCA TCT TCA TCC CCA AAC GCT GAC ATG GAG GCC CAG AAT AC |
| KRE6_del_PR2 | *KRE6a*/*b* and *KRE6b* deletion cassette construction | GCA AAA CCA AGC GAA GGA ATA GCC AGT ATA GCG ACC AGC ATT CAC |
| KRE6_del_PF3 | *KRE6a*/*b* and *KRE6b* deletion cassette construction | GTG AAT GCT GGT CGC TAT ACT GGC TAT TCC TTC GCT TGG TTT TGC |
| KRE6_del_PR3 | *KRE6a*/*b* and *KRE6b* deletion cassette construction | TTA CAT GGC CAT GAA AAT GGC GC |
| KRE6_del_PF4 | *KRE6a*/*b* and *KRE6a* deletion cassette construction | AAA ATC TGA GGC TGT GTG TCG C |
| KRE6_del_PR4 | *KRE6a*/*b* and *KRE6b* deletion cassette construction | AAG TCG ATC CGA GTC AGG TG |
| KRE6a_del_PR2 | *KRE6a* deletion cassette construction | CCA CAA CGT CAA GTT GGG GTC AGT ATA GCG ACC AGC ATT CAC |
| KRE6a_del_PF3 | *KRE6a* deletion cassette construction | GTG AAT GCT GGT CGC TAT ACT GAC CCC AAC TTG ACG TTG TGG |
| KRE6a_del_PR3 | *KRE6a* deletion cassette construction | TCC CGA TCC ATG CTA CCT TG |
| KRE6a_del_PR4 | *KRE6a* deletion cassette construction | CCT TCT ACT TCT CTG CCT CTC |
| KRE6b_del_PF1 | *KRE6b* deletion cassette construction | CCA TCA CCA TCA CCT TCA ATG C |
| KRE6b_del_PR1 | *KRE6b* deletion cassette construction | GTA TTC TGG GCC TCC ATG TCA GCG AGC AGC TAT GAG GAA AAA G |
| KRE6b_del_PF2 | *KRE6b* deletion cassette construction | CTT TTT CCT CAT AGC TGC TCG CTG ACA TGG AGG CCC AGA ATA C |
| KRE6b_del_PF4 | *KRE6b* deletion cassette construction | TTG TTT TTG TGG CGC CAG CC |
| KRE6_del_verif_PF | *KRE6a*/*b* deletion verification | AGG CTT AGT GAG AAA CCC CTA C |
| NatR_verif_PR | *KRE6a*/*b* deletion and *KRE6b* deletion verification | AGC ATC ACC TGG AAC AGA AGT TC |
| KRE6_PF | *KRE6a*/*b* deletion verification | AGA TAG CGC CCA TGG ACA TC |
| KRE6_PR | *KRE6a*/*b* deletion and *KRE6a* deletion verification | TAG TTT CAG ACG ACC CAC CTC |
| NAT1_743_PF | *KRE6a* deletion verification | GTG CTG GTC ATT TGT GGT TG |
| KRE6a_PR | *KRE6a* deletion verification | ATG ATG CCT TGG CTT CGA CTG |
| KRE6a_PF4 | *KRE6b* deletion verification | ATC CAG CAA GCT GTT TCG GG |
| KRE6b_PF2 | *KRE6b* deletion verification | GAG GTG GGT CGT CTG AAA CTA |
| KRE6b_PR(sybrgreen) | *KRE6b* deletion verification | CTT GAA CCC TGC CAT CTC CC |
| KRE6_del_sg5’ | *KRE6a*/*b* deletion and *KRE6a* guide RNA | ACU GUG UUC UGG GAC UGU GG |
| KRE6_del_sg3’ | *KRE6a*/*b* deletion and *KRE6b* deletion guide RNA | AUG CCG AAG AAC AAA CUC AG |
| ***KRE6a*/*b* overexpression** | | |
| KRE6a_PF_KasI | *KRE6a* overexpression plasmid construction | ACA CTG GCG CCA TGG TCC GTG ACT TGA CCT C |
| KRE6a_PR_NruI | *KRE6a* overexpression plasmid construction | ACA CTT CGC GAT CAA CAG TCG TAG GCC AAC TTG |
| KRE6b_PF_KasI | *KRE6b* overexpression plasmid construction | ACA CTG GCG CCA TGT CCC ACA GAG ACC TCA C |
| KRE6b_PR_NruI | *KRE6b* overexpression plasmid construction | ACA CTT CGC GAC TAG CAA CCA CTG AGT TTG TTC TT |
| pjli8_ADH1_PF | *KRE6a*/b overexpression plasmid sequencing | AGC AAC ACC GGT GGA ATT TCC |
| KRE6a_PF2 | *KRE6a* overexpression plasmid sequencing | TTG CAC AAC CCA GAC CCA ATT G |
| KRE6a_PF3 | *KRE6a* overexpression plasmid sequencing | GCC ACA ATT TGT TCT ACC GCT C |
| KRE6a_PF4 | *KRE6a* overexpression plasmid sequencing | ATC CAG CAA GCT GTT TCG GG |
| KRE6b_PF2 | *KRE6b* overexpression plasmid sequencing | GAG GTG GGT CGT CTG AAA CTA |
| KRE6b_PF3 | *KRE6b* overexpression plasmid sequencing | GAA CTT TCT ACG ATG GCG ACG |
| KRE6b_PF4 | *KRE6b* overexpression plasmid sequencing | GAC ACT CTC AAA ACT GGT GTG G |
| CauNi_sg5’ | *KRE6a*/*b* overexpression guide RNA | CCC GGA GAU ACA CGG CGC CG |
| CauNi_sg3’ | *KRE6a*/*b* overexpression guide RNA | GCU GCA AAA UAA GGC CAG AG |
| **RT-PCR** | | |
| ACT1_F(sybrgreen) | RT-PCR | GAA GGA GAT CAC TGC TTT AGC C |
| ACT1_R(sybrgreen) | RT-PCR | GAG CCA CCA ATC CAC ACA G |
| KRE6a_PF(sybrgreen) | *KRE6a* RT-PCR | TGA CGT GGT ATG TGG GAA GC |
| KRE6a_PR(sybrgreen) | *KRE6a* RT-PCR | GGG CTC TTT GGA AAT GCG TC |
| KRE6b_PF(sybrgreen) | *KRE6b* RT-PCR | TCC TGA AGA CTA CCC GAC GT |
| KRE6b_PR(sybrgreen) | *KRE6b* RT-PCR | CTT GAA CCC TGC CAT CTC CC |
| FKS1_PF(sybrgreen) | *FKS1* RT-PCR | GTA TGG GTT ACA TGG CCG CT |
| FKS1_PR(sybrgreen) | *FKS1* RT-PCR | GCA AGA ATG AAG TCA CCG GC |
| CHS1_PF(sybrgreen) | *CHS1* RT-PCR | CCA GGA GAA ACG GGC AGA AA |
| CHS1_PR(sybrgreen) | *CHS1* RT-PCR | TAA CCC GTA GAG CCA ATC GC |

**Supplementary Table 9. ^13^C and ^15^N Solid-state NMR experimental parameters for fungal cell wall characterization.** T = sample temperature; B_0_ = magnetic field; ν_MAS_ = MAS frequency; ns = number of scans; d_1_ = recycle delay between scans; t_1, max_ = maximum t_1_ evolution time (for indirect dimension); t_1, inc_ = increment for t_1_ (for indirect dimension) evolution time; τ_dw_ = dwell time during direct FID acquisition; τ_acq_ = maximum acquisition time during direct FID detection; τ_XY_ = cross-polarization contact time during CP from channel X to channel Y; ν_1H, dec_ = dipolar decoupling field strength**.** DNP experiments are marked with asterisks. Spin diffusion (SD).

| **Experiment** | **NMR Parameters** | | | | | | | | | | | | | | | **Samples** |
| --- | --- | --- | --- | --- | --- | --- | --- | --- | --- | --- | --- | --- | --- | --- | --- | --- |
|  | T  (K) | B_0_ (T) | ν_MAS_ (kHz) | ns | d1  (s) | t_1, max_ (ms) | t_1, inc_ (μs) | τ_dw_ (μs) | τ_acq_ (ms) | τ_HC_ (ms) | τ_HN_ (ms) | τ_NC_ (ms) | τ_SD_ (ms) | τ_mix_ (ms) | ν_1H dec_ (kHz) |  |
| **Identification and quantification of polysaccharides** | | | | | | | | | | | | | | | |  |
| 1D ^13^C CP | 298 | 18.8 | 13.5 | 256 | 1.8 |  |  | 7 | 18 | 1 |  |  |  |  | 83 | C. al apo  C.al CAS  (SC5314) C. au apo  C.au CAS  (AR386,  P*_ADH1__KRE6a*) |
| 1D ^15^N CP | 298 | 18.8 | 13.5 | 128 | 2 |  |  | 10 | 16 |  | 1 |  |  |  | 83 |  |
| 1D ^13^C DP | 298 | 18.8 | 13.5 | 32-64 | 2 or 30 |  |  | 7 | 29 |  |  |  |  |  | 100 |  |
| 1D ^13^C refocused INEPT | 298 | 18.8 | 13.5 | 256 | 4 |  |  | 7 | 29 |  |  |  |  | 1.7  τ_J_ | 71 |  |
| 2D ^13^C-^13^C with CORD mixing | 298 | 18.8 | 13.5 | 16 | 1.5 | 7 | 29 | 7.5 | 18 | 0.5 |  |  |  | 53 τ_CORD_ | 83 |  |
| 2D ^13^C-^13^C refocused DP J-INADEQUATE | 298 | 18.8 | 13.5 | 8 | 1.5 | 10 | 20 | 7.5 | 19 |  |  |  |  |  | 83 |  |
| 2D ^15^N-^13^C N(CA)CX with DARR mixing | 298 | 18.8 | 13.5 | 64 | 1.6 | 7 | 180 | 7.5 | 18 |  | 0.6 | 5 |  | 100 τ_DARR_ | 93 |  |
| 2D ^1^H-^13^C refocused INEPT | 298 | 18.8 | 13.5 | 4 | 2 | 11 | 50 | 7.5 | 23 |  |  |  |  | 1.7  τ_J_ | 71 |  |
| 2D ^1^H-^15^N HETCOR with SD | 298 | 18.8 | 13.5 | 16 | 1.6 | 2 | 41 | 7.5 | 18 |  | 1 |  | 1 |  | 93 |  |
| **Estimation of site-specific hydration of polysaccharides** | | | | | | | | | | | | | | | |  |
| 2D ^13^C-^13^C water-edited | 290 | 9.4 | 10 | 128- 256 | 1.6 | 5 | 71 | 10 | 14 | 1 |  |  | 0, 4 | 50  τ_PDSD_ | 71 |  |
| **Dynamics of polysaccharides** | | | | | | | | | | | | | | | |  |
| 1D ^13^C-T_1_ | 298 | 9.4 | 10 | 128- 256 | 2 |  |  | 10 | 16 | 1 |  |  |  |  | 71 |  |
| 1D ^1^H-T_1ρ_ | 298 | 9.4 | 10 | 256 | 2 |  |  | 10 | 14 | 1 |  |  |  |  | 71 |  |
| **Intermolecular interactions of polysaccharides** | | | | | | | | | | | | | | | |  |
| * 2D ^13^C-^13^C with PAR mixing | 92 | 14.1 | 8 | 4 | 5 | 7 | 33 | 7.5 | 15 | 0.5 |  |  |  | 5-20  τ_PAR_ | 93 | C. au apo  C.au CAS |

**Supplementary Table 10.  Parameters used for proton detection experiments.** The CP based 2D hCH proton detection experiments were performed on 600 MHz (14.1 T) spectrometer with the MAS frequency of 60 kHz and 2D and 3D hCCH TOCSY (DIPSI-3) was performed on 800 MHz (18.8 T) with the MAS frequency of 13.5 kHz.

| **Expt.** | **Samples** | **Temperature**  **(K)** | **CP (µs)** | | **D1** | **NS** | **td2** | **td1** | **td3** | **aq2**  **(ms)** | **aq1**  **(ms)** | **aq3**  **(ms)** | **Decoupling** | **Water suppression** | ***J-*evolution**  **(ms)** | **DIPSI-3**  **(ms)** |
| --- | --- | --- | --- | --- | --- | --- | --- | --- | --- | --- | --- | --- | --- | --- | --- | --- |
|  |  |  | t_cp1_ | t_cp2_ |  |  |  |  |  |  |  |  |  |  |  |  |
| 2D hCH | *C. albicans*  *C.auris* | 304  300 | 900 | 100 | 2 | 64 | 1764 | 320 | - | 14.9 | 5.3 | - | slpTPPM  (rf 20.280 kHz) | MISSISSIPI  (total duration)  100 ms  (rf 30 kHz) | - | - |
| 2D & 3D hCCH TOCSY (DIPSI-3) | *C. albicans*  *C.auris* | 296  295 | - | - | 1.89 | 8 | 2614 | 128 | 128 | 39.9 | 2.56 | 2.56 | SPINAL-64  (rf 71.429 kHz)  WALTZ-16  (rf 10 kHz) | MISSISSIPI  (total duration)  40 ms  (rf 25.994 kHz | 1.78 (τ_1_)  1.19 (τ_2_ | 25.5 |
